# Small molecule modulator of neuronal lysosome positioning and function resolves Alzheimer’s Disease-linked pathologies in cultured human neurons

**DOI:** 10.1101/2024.11.04.621986

**Authors:** Amanda M. Snead, Sruchi Patel, Mia Krout, Ryan S. Hippman, Gabrielle Smith, Diya Dileep, Nitya Chagoor, Rachel Shi, Ricardo Linares, Andrew Dobria, Stephanie M. Cologna, Camerron Crowder, Leslie N. Aldrich, Swetha Gowrishankar

**Author notes:** contributed equally (Diya Dileep, Nitya Chagoor). contributed equally (Rachel Shi, Ricardo Linares).

## Abstract

Abnormal increase in axonal lysosome abundance is associated with multiple neurodegenerative diseases including Alzheimer’s disease. However, the underlying mechanisms and disease relevance are not fully understood. We have recently identified RH1115 as a small molecule modulator of the autophagy-lysosomal pathway that regulates lysosome positioning in neurons. This allowed us to manipulate neuronal lysosome distribution including in axons and interrogate its contribution to both optimal neuronal functioning and to disease pathology. We demonstrate that the small molecule not only rescues aberrant buildup of both axonal autophagic and lysosomal intermediates but also reduces secreted Aβ42 levels in human iPSC-derived neurons lacking the lysosomal adaptor, JIP3. We thus demonstrate that restoring efficient axonal lysosome transport has an anti-amyloidogenic effect in human neurons and is a promising therapeutic strategy for Alzheimer’s disease. Furthermore, we show that the small molecule enhances neuronal lysosome degradation, requires the lysosomal adaptor JIP4 to rescue the axonal lysosome pathology in the JIP3 KO neurons and increases levels of the JIP4-interacting lysosomal membrane protein, TMEM55B. Lastly, treatment with the small molecule led to a striking rescue of locomotor defects in JIP3 KO zebrafish larvae. Thus, we have identified a small molecule which can be impactful in neurodegenerative diseases that have a lysosomal pathology and have determined its molecular targets in modulating axonal lysosome abundance.

## Introduction

Lysosomes are the central organelles in maintaining protein and organelle homeostasis in neurons over their long lifespan. In addition to their longevity, their polarity and extreme size add additional spatial challenges for optimal lysosome biogenesis and functioning in neurons. This necessitates efficient lysosome transport as an important adaptation to meet the metabolic demands of the neuron. Indeed, efficient axonal lysosome transport is coupled to the maturation of these organelles and necessary for clearance of cargo from axons (Ferguson, 2018; Paumier & Gowrishankar, 2024; Sidibe et al., 2022). Multiple studies suggest that autophagosomes, which play an important role in clearing damaged organelles and old proteins from axons, engage with endolysosomes and undergo retrograde transport so that cargo can be degraded in the highly acidic lysosomes in the soma (Cheng et al., 2015a; Kuijpers et al., 2021; Maday et al., 2012; Maday & Holzbaur, 2014). The transport of lysosomes and their precursor organelles along the axon requires the modulation of microtubule tracks by post-translational modification, microtubule associated proteins (MAPs), molecular motors, and adaptor proteins that regulate the connections between motors and their cargo (Berth & Lloyd, 2023; Ferguson, 2018; Fu & Holzbaur, 2014; Paumier & Gowrishankar, 2024).

The importance of efficient axonal lysosome transport to maintenance of neuronal health and functioning is exemplified by how disruptions to this process are linked to several neurological diseases (Berth & Lloyd, 2023; Ferguson, 2018; Paumier & Gowrishankar, 2024; Roney et al., 2022; Xiong & Sheng, 2024). A high abundance of axonal lysosomes within swellings at sites of amyloid plaques have been observed in Alzheimer’s disease (Cataldo et al., 1996; Condello et al., 2011; Gowrishankar et al., 2015; Nixon et al., 2005; Terry et al., 1964; Tsai et al., 2004). These organelles have been linked to increased Amyloid Precursor Protein (APP) processing and exacerbated plaque development in AD (Gowrishankar et al., 2015, 2017; Kandalepas et al., 2013; Tammineni & Cai, 2017). Indeed, loss of JNK-interacting protein 3 (JIP3), a brain-enriched adaptor (Kelkar et al., 2003) and a critical regulator of axonal lysosome transport (Drerup & Nechiporuk, 2013; Edwards et al., 2013; Gowrishankar et al., 2017, 2021), results in a similar focal buildup of axonal lysosomes within swellings as well as increased intraneuronal amyloid β42 (Aβ42) levels (Gowrishankar et al., 2017). Furthermore, haploinsufficiency of JIP3 in an AD mouse model resulted in worsening of amyloid pathology (Gowrishankar et al., 2017), supporting a model wherein these stalled axonal organelles lead to increased amyloid processing and pathology development. Thus, restoring efficient axonal lysosome transport and maturation could suppress amyloidogenic processing of APP and in turn, the progression and development of AD pathology. To this end, we now show that a small molecule modulator of <u>a</u>utophagic and lysosomal <u>p</u>athways (ALP) that our two groups recently identified (Hippman et al., 2023), RH1115, can rescue both the axonal lysosome accumulation pathology as well as the aberrant Aβ42 production found in JIP3 KO human iPSC-derived neurons (i^3^Neurons). We show that RH1115 can also reduce the buildup of autophagic vacuoles within neuronal processes of JIP3 KO i^3^Neurons suggesting that it acts to enhance retrograde transport of these axonal organelles, and hence their maturation. We find that this rescue of axonal lysosome buildup in JIP3 KO i^3^Neurons by RH1115 requires JIP4, an adaptor highly related to JIP3, that has been implicated in regulating dynein-based retrograde lysosome movement in cells (Sasazawa et al., 2022; Vilela et al., 2019; Willett et al., 2017). RH1115 also leads to increased levels of the lysosomal transmembrane protein TMEM55B, a JIP4-interacting protein shown to recruit it to lysosomes and mediate starvation-induced lysosome movement in cultured cells (Willett et al., 2017). Our study provides a proof of principle that restoring normal axonal lysosome transport and clearing organelles that build up there, can modulate amyloid production and is therefore a viable therapeutic avenue to pursue. In further support of enhancement of axonal lysosome transport efficiency as a strategy to restore optimal neuronal health and function, we show that the small molecule rescued locomotor defects in JIP3 KO zebrafish larvae.

In addition to its effect on axonal lysosomes, RH1115 mobilized lysosomes within the neuronal cell body to a more perinuclear location and enhanced their degradative capacity. Lysosome positioning and motility within cells is regulated by several factors including nutrient status (Ballabio & Bonifacino, 2020a; Pu et al., 2016). In turn, lysosome positioning can affect different cellular functions including signaling, autophagy, cell migration and adhesion (Ballabio & Bonifacino, 2020; Korolchuk et al., 2011). Lysosomes differ in their intraluminal pH and degradative capacity depending on their positioning within the cells, with peripheral lysosomes being less degradative and perinuclear lysosomes being more degradative and conducive to receiving autophagic cargo for turnover (Gowrishankar & Ferguson, 2016; Johnson et al., 2016). Given the ability of the small molecule to mobilize lysosomes to the perinuclear region and its ability to enhance lysosome function, it may be relevant to the treatment of other diseases characterized by altered lysosome function and distribution including lysosomal storage diseases.

## Results

### Loss of JIP3 in human i^3^Neurons reduces motility of both autophagosomes and autolysosomes, but not fusion of autophagosomes with endo-lysosomes

Mammalian JIP3/ MAPK8IP3 as well as its orthologs in the nematode worm and zebrafish have been shown to play a critical role in axonal lysosome transport with their loss resulting in an abnormal accumulation of endo-lysosomes within axonal swellings (Drerup & Nechiporuk, 2013; Edwards et al., 2013; Gowrishankar et al., 2017, 2021). Autophagosome biogenesis and trafficking in axons is spatially organized, and dependent on fusion with endo-lysosomes for acquiring adaptors that aid in their retrograde transport to the soma for degradation (Cheng et al., 2015a; Maday et al., 2012; Maday & Holzbaur, 2014). Given the critical role of the lysosome adaptor JIP3 in regulating retrograde lysosome transport, we predicted that its loss would affect the movement and distribution of autophagic vacuoles in neurons lacking JIP3. Towards this end, we examined the dynamics of LC3-containing vesicles in the neurites of Control and JIP3 KO i^3^Neurons generated from iPSCs stably expressing LC3-RFP-GFP. We found that there were far more autophagic vacuoles in the neurites of JIP3 KO i^3^Neurons, both within axonal swellings (Figure 1 A; white arrowheads) and outside of the swellings, compared with the neurites of Control i^3^Neurons (Figure 1 A). Furthermore, our timelapse imaging studies revealed a significant decrease in the motile fraction of both autophagosomes (both RFP- and GFP-positive) and the more acidic autolysosomes (only RFP-positive) (Figure 1 B, C) in JIP3 KO i^3^Neurons compared to Control i^3^Neurons. We next examined if loss of JIP3 had any effect on fusion of autophagosomes with the endolysosomes in the neurites. Interestingly, we found that the percentage of autolysosomes in the neurites of JIP3 KO i^3^Neurons was comparable to that in Control i^3^Neurons (Figure 1 D), suggesting that JIP3 is not necessary for the fusion of autophagosomes with endo-lysosomes in these neuronal processes.

**Figure 1:**
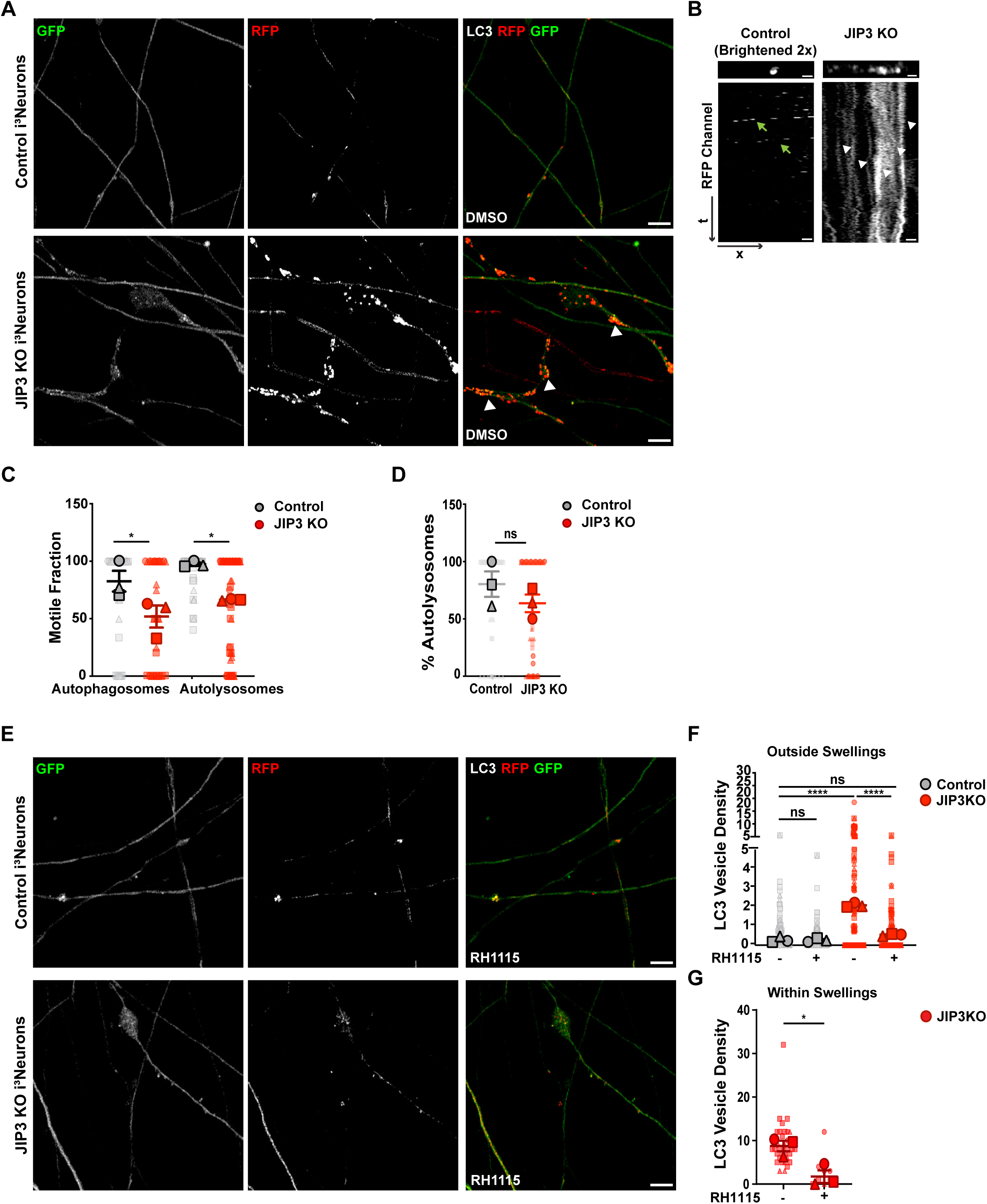
Loss of JIP3 impairs motility of autophagosomes and autolysosomes, causing their buildup in neuronal processes, which is rescued by small molecule activator of ALP. **A)** Individual gray scale and merged images at t=10 from time lapse Airyscan images acquired at 2.5 frames/sec of DIV15-16 Control and JIP3 KO i^3^Neurons stably expressing LC3-RFP-GFP treated with 0.15% DMSO for 72 h. White arrowheads point to axonal swellings filled with autophagosomes and autolysosomes in JIP3 KO i^3^Neurons. Scale bar, 5µm. **B)** Portion of straightened representative neurite (top) and corresponding kymographs (bottom) depicting LC3-RFP-GFP (red channel) vesicle density and movement, respectively, in Control and JIP3 KO i^3^Neurons. Scale bar, 1µm. Green arrows point to moving vesicles, white arrowheads point to stationary vesicles. Control images are shown 2× brighter than JIP3 KO to enhance their visibility. **C)** Quantification depicting the motile fraction of autophagosomes (RFP+ and GFP+) and autolysosomes (RFP+ only) in DIV15-16 Control and JIP3 KO i^3^Neurons. Superplots show mean ± SEM as well as individual data (represented as large and small symbols respectively. Circles, triangles, and squares represent each independent experiment). N=3 independent experiments, Control autolysosomes n=99 neurites, JIP3 KO autolysosome n=69 neurites, Control autophagosome n=47 neurites, JIP3 KO autophagosome n=40 neurites; \**p*<.05; one-way ANOVA with Sidak’s multiple comparisons. **D)** Quantification showing percentage of total autophagic vacuoles that are autolysosomes in DIV15-16 Control and JIP3 KO i^3^Neurons. Superplots show mean ± SEM as well as individual data (represented as large and small circles, triangles and squares). N=3 independent experiments, Control n=45 vesicles, JIP3 KO n=58 vesicles; ns=not significant; unpaired t-test. **E)** Individual gray scale and merged snapshots at t=10 from time lapse Airyscan images acquired at 2.5 frames/sec of DIV15-16 Control and JIP3 KO i^3^Neurons stably expressing LC3-RFP-GFP and treated for 72 h with 15µM RH1115. Scale bar, 5µm. **F)** Quantification of autophagic vacuole density (LC3 vesicle number per 10µm of neurite) outside of swellings. Superplots show mean ± SEM as well as individual data (represented as large and small circles, triangles and squares). N=3 independent experiments. Control DMSO n=120 neurites, Control RH1115 n=125 neurites, JIP3 KO DMSO n=141 neurites, JIP3 KO RH1115 n=130 neurites; \*\*\*\**p*<.0001; one-way ANOVA with Sidak’s multiple comparisons. **G)** Quantification of autophagic vacuole density within swellings. Superplots show mean ± SEM as well as individual data (represented as large and small symbols). N=3 independent experiments, DMSO n=33 swellings, RH1115 n=9 swellings in JIP3 KO i^3^Neurons (only found in JIP3 KO genotype) \**p*<.05; unpaired t-test.

### RH1115 clears autophagic vacuole buildup from neuronal processes of JIP3 KO i^3^Neurons

In collaboration with the Aldrich lab, we have recently identified a small molecule modulator of the ALP, RH1115, that enhances autophagic flux as well as alters lysosome positioning in both HeLa cells and the cell bodies of i^3^Neurons (Hippman et al., 2023). Since RH1115 strongly increased perinuclear localization of lysosomes in Control i^3^Neurons, suggestive of increased net retrograde lysosome movement, we tested whether the small molecule could mobilize and effect retrograde movement of the autophagic vacuoles that build up in the neuronal processes of JIP3 KO i^3^Neurons. We found that RH1115 treatment led to a dramatic decrease in the autophagic vacuole accumulation observed in JIP3 KO i^3^Neurons (Figure 1 A, E, F). There is an almost four-fold increase in autophagic vacuole density in neuronal processes of JIP3 KO i^3^Neurons, even outside of axonal swellings, which is rescued to levels comparable to that of Control i^3^Neurons (Figure 1 F). Importantly, the small molecule can reduce even the massive autophagic vacuole build up observed within the axonal swellings of JIP3 KO i^3^Neurons (Figure 1 A, E, G).

### RH1115 reduces both axonal lysosome build up and Aβ42 pathology of JIP3 KO i^3^Neurons and increases net retrograde movement of the lysosomes in the soma

Given the strong reduction of autophagic vacuole buildup effected by RH1115 in JIP3 KO i^3^Neurons, and their previously established dependence on axonal lysosomes for their transport to the soma for clearance (Cheng et al., 2015b; Maday et al., 2012; Maday & Holzbaur, 2014), we next examined how RH1115 affected axonal lysosome pathology in these i^3^Neurons. As previously shown (Gowrishankar et al., 2021), JIP3 KO i^3^Neurons exhibit a strong axonal lysosome accumulation phenotype which is both robust and penetrant (Figure 2 A; S1A, B). RH1115 treatment strongly reduced the axonal lysosome buildup (Figure 2 A, B; S1A, B). In fact, the small molecule almost eliminated (97% reduction) the larger lysosome accumulations that would often extend well over 10 microns in length in these JIP3 KO i^3^Neurons (Figure 2 A, B). In addition, RH1115 strongly suppressed the overall axonal lysosome buildup (Figure 2 A, B), leading to a lysosome distribution like that observed in Control i^3^Neurons (Figure 2 A). We also observed a similar rescue of axonal lysosome buildup in JIP3 KO i^3^Neurons stably expressing LAMP1-GFP upon treatment with RH1115 (S1 A, B).

**Figure 2:**
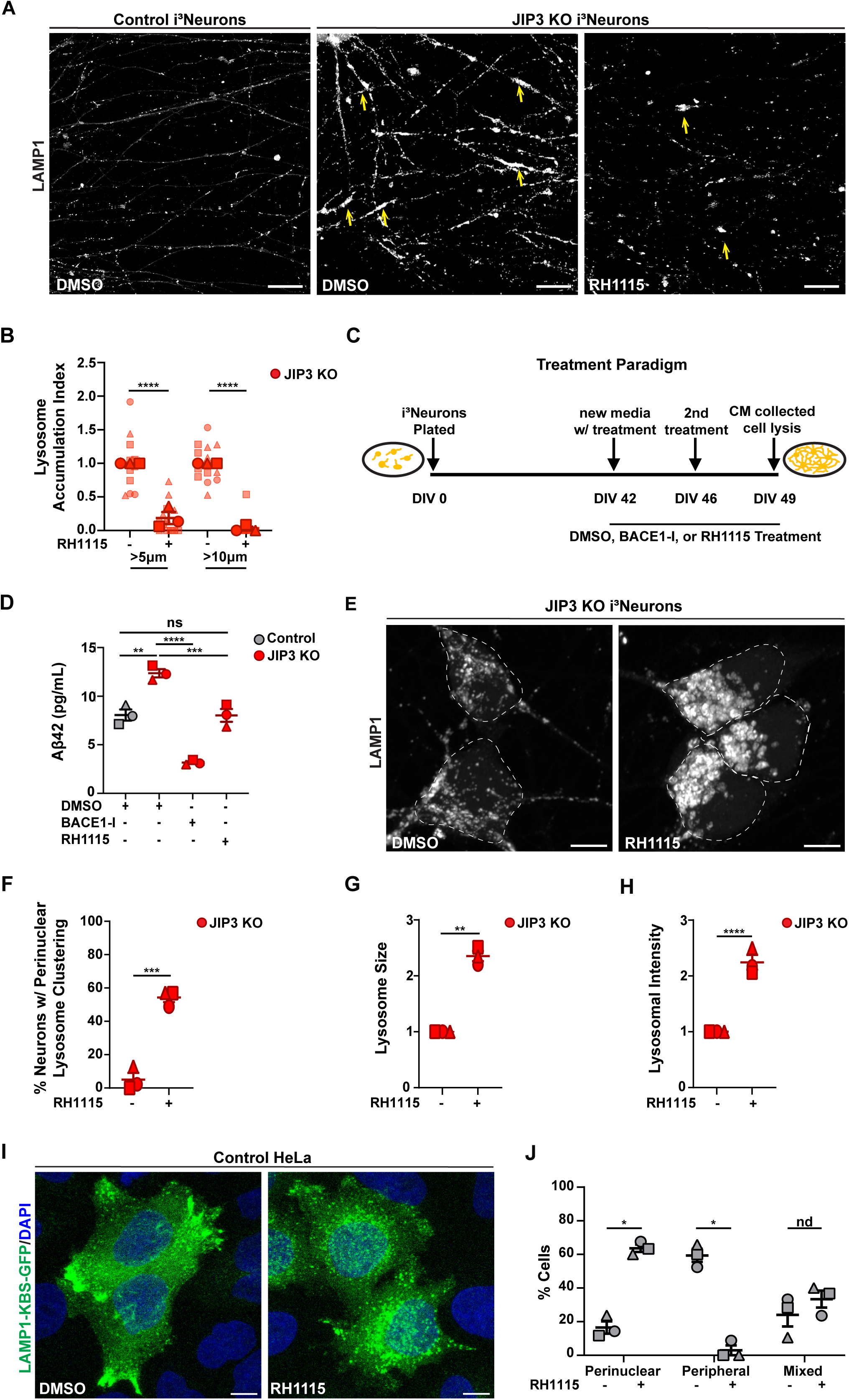
RH1115 reduces the axonal lysosome buildup observed in JIP3 KO i^3^Neurons and increases perinuclear lysosome clustering in neuronal cell bodies of these i^3^Neurons. **A)** High resolution, stitched, greyscale images of neuronal processes of DIV11 Control and JIP3 KO i^3^Neurons stained for endogenous LAMP1 after 72 h treatment with 0.15% DMSO or 15μM RH1115. Arrows point to LAMP1-positive vesicle accumulations. Scale bar, 20μm. **B)** Quantification of the axonal lysosome accumulation index (swellings greater than 5μm or greater than 10μm normalized to neurite density). Superplots show normalized mean ± SEM as well as individual data (represented as large and small circles, triangles and squares). N=3 independent experiments, n=14 stitched images per condition. \*\*\*\**p*<.0001; one-way ANOVA followed by Sidak’s multiple comparisons test. **C)** Treatment paradigm for Aβ42 measurement: i^3^Neurons were plated on DIV0 and differentiated until DIV42/43 when media was changed for a weeklong drug treatment. i^3^Neurons were treated with 0.15% DMSO, 15μM RH1115 or 5μM BACE1-I two times over the week (DIV42/43 and DIV47/48) and the conditioned media was collected when the i^3^Neurons were lysed on DIV49/50. **D)** Extracellular Aβ42 was measured by ELISA using the media collected from DIV49/50 Control and JIP3 KO i^3^Neurons following the treatment paradigm described above. One-way ANOVA followed by Sidak’s multiple comparisons. \*\**p*<.01, \*\*\**p*<0.001, \*\*\*\**p*<0.0001, ns=not significant. Different symbol shapes represent 3 independent replicates. **E)** High resolution greyscale images of DIV11 JIP3 KO i^3^Neuron somas treated with 0.15% DMSO or 15μM RH1115 for 72 h and stained for endogenous LAMP1. Somas are outlined with a white dashed line. Scale bar, 5μm. **F)** Quantification of the percent of DIV10 JIP3 KO i^3^Neurons with LAMP1 perinuclear localization, plots show mean ± SEM, N=3 independent experiments, DMSO treated n=106, RH1115 n=123, \*\*\**p* <.001; unpaired t-test. **G)** Quantification of the average LAMP1-vesicle size in DIV10 JIP3 KO i^3^Neurons, plots show mean ± SEM, N=3, DMSO n=21 somas, RH1115 n=22 somas, \*\**p*<.01; unpaired t-test. **H)** Quantification of the average endogenous LAMP1 fluorescence intensity in DIV10 JIP3 KO i^3^Neurons, plots show mean ± SEM, N=3 independent experiments, DMSO n=21 somas, RH1115 n=22 somas, \*\*\*\**p*<.0001; unpaired t-test. **I)** Confocal images of Control HeLa cells transfected with LAMP1-KBS-GFP and treated with 0.15% DMSO or 15μM RH1115 for 6 h, 48h post-transfection. Cells were stained with anti-GFP and DAPI, Scale bar, 10μm. **J)** Quantification of the percent of cells showing the majority of LAMP1-KBS-GFP in the cell periphery, perinuclear area, or a mixed distribution. Plots show mean ± SEM, N=3 independent experiments (represented as circles, triangles, and squares). DMSO n=91 cells, RH1115 n=84 cells; \**p*<0.05, nd=no discovery; grouped analysis multiple unpaired t-tests. P value of individual t-tests for peripheral and perinuclear localization, \*\*\**p*<0.001.

In addition to the axonal lysosome phenotype, JIP3 KO i^3^Neurons have previously been shown to have higher levels of intraneuronal Amyloid β42 (Aβ42) (Gowrishankar et al., 2021). We found that JIP3 KO i^3^Neurons in fact also have higher levels (50% increase) of extracellular or secreted Aβ42, as measured from the culture media (Figure 2 C, D). Given links between altered axonal lysosome transport, maturation and increased amyloidogenesis (Adalbert et al., 2009; Gowrishankar et al., 2015, 2017, 2021; Kandalepas et al., 2013; Orlowski et al., 2024; Tammineni & Cai, 2017), and the dramatic rescue of axonal lysosome pathology by RH1115, we examined if the small molecule had any effect on Aβ42 production in these JIP3 KO i^3^Neurons. Indeed, we found that RH1115 treatment significantly decreased extracellular Aβ42 in JIP3 KO i^3^Neurons, to levels comparable to Control i^3^Neurons (Figure 2 C, D). Importantly, the small molecule had no negative impact on neuronal viability at the dosage at which it rescues axonal lysosome buildup in JIP3 KO i^3^Neurons (Figure S1C).

Given the clearance of axonal lysosome buildup in JIP3 KO i^3^Neurons brought about by RH1115, we next examined how it modulated movement as well as properties of lysosomes in the neuronal cell body. We found that RH1115 causes a strong perinuclear localization of lysosomes within the soma (Figure 2 E, F; S1D), as previously observed in Control i^3^Neurons (Hippman et al., 2023). We also observe an increase in the size and intensity of LAMP1-positive vesicles, in the soma of JIP3 KO i^3^Neurons (Figure 2 E, G, H). Based on the mobilization and clearance of axonal lysosomes and the perinuclear clustering of soma lysosomes induced by RH1115, we hypothesized that the small molecule causes an increase in the net retrograde movement of lysosomes. We tested this by examining the effect of RH1115 on distribution of LAMP1-KBS-GFP expressed in HeLa cells. This construct includes LAMP1 with three copies of the Kinesin light chain binding sequence of SKIP, tagged with GFP (Pernigo et al., 2013; Pu et al., 2015). This bypasses the requirement for ARL8B, SKIP and BORC, allowing for direct binding between LAMP1 and kinesin-1, which results in a strong peripheral accumulation of lysosomes (Pu et al., 2015). As expected, a little over 60% of DMSO-treated LAMP1-KBS-GFP expressing cells show a strong peripheral LAMP1-KBS-GFP accumulation, with only about 20% cells showing a perinuclear LAMP1 distribution (Figure 2 I, J). However, in the case of RH1115 treatment, there is a dramatic shift in lysosome positioning in most cells, with greater than 60% of cells showing LAMP1-KBS-GFP strongly enriched in the perinuclear region, and almost none having lysosomes mainly in the periphery (Figure 2 I, J).

### RH1115 increases neuronal lysosomal degradative capacity

Lysosome positioning in non-neuronal cells has been linked to functional differences, with lysosomes that are more perinuclear being more degradative than peripheral lysosomes (Gowrishankar & Ferguson, 2016; Johnson et al., 2016). Given the effect of RH1115 in mobilizing neuronal lysosomes towards the perinuclear region, we examined its impact on lysosome acidification. Control and JIP3 KO i^3^Neurons treated with RH1115 for 72 hours were labelled with Lysotracker Red, a fluorescent dye that preferentially identifies acidic organelles. RH1115 treatment led to a ten-fold increase in intensity of Lysotracker Red in both Control and JIP3 KO i^3^Neurons (Figure 3 A, B), suggestive of an increase in acidic organelles in the neuronal cell body.

**Figure 3:**
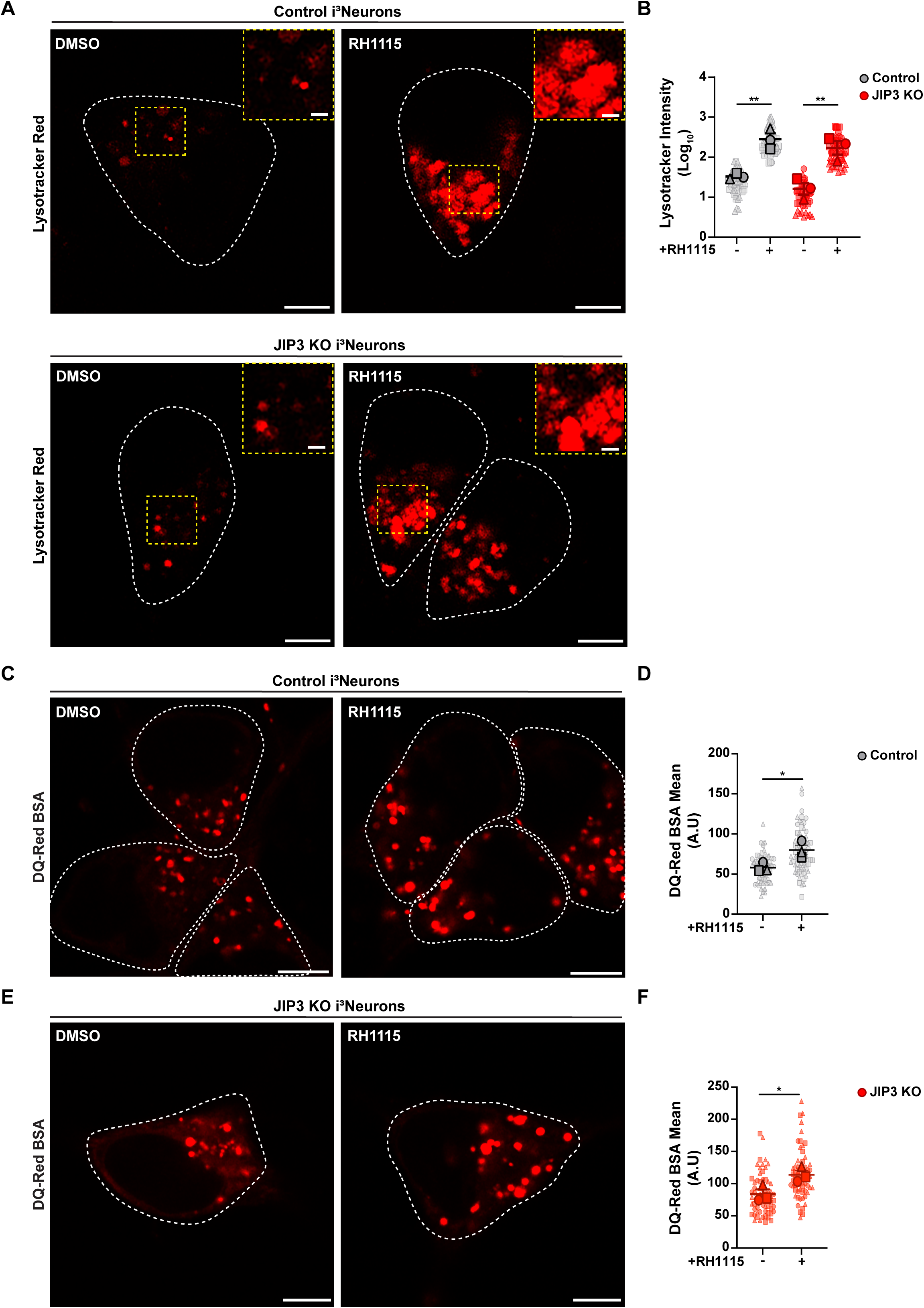
RH1115 treatment results in increased acidic lysosomes and enhanced lysosomal degradation in both Control and JIP3 KO i^3^Neurons. **A)** Confocal images of DIV10-13 Control and JIP3 KO i^3^Neurons treated with 0.15% DMSO or 15µM RH1115 for 72 h and stained for Lysotracker Red for 10 minutes. Scale bar, 5µm. Neuronal cell bodies are outlined with a dashed line. Insets show higher magnification image of area outlined by the dashed box within the image. Scale bar, 1µm. **B)** Quantification of the mean lysotracker intensity per soma in DIV10-13 Control and JIP3 KO i^3^Neurons, transformed using Log_10_ to account for the exponential difference between DMSO and RH1115 fluorescence intensity values. Superplots show mean ± SEM as well as individual data (represented as large and small circles, triangles and squares). N=3 independent experiments, Control DMSO n=47 i^3^Neurons, Control RH1115 n=43 i^3^Neurons, JIP3 KO DMSO n=37 i^3^Neurons, JIP3 KO RH1115 n=55 i^3^Neurons; \*\**p*<.01; one way ANOVA followed by Sidak’s multiple comparisons. **C)** Confocal images of DIV12-13 Control i^3^Neuron somas pre-loaded with 25µg/mL DQ-Red BSA and treated with 0.15% DMSO or 15µM RH1115. Scale bar, 5µm. **D)** Quantification of the mean intensity of DQ-Red BSA. Superplots show mean ± SEM as well as individual data (represented as large and small circles, triangles and squares). N=3 independent experiments, Control DMSO n=72 i^3^Neurons, Control RH1115 n=71 i^3^Neurons, \**p*<.05; unpaired t-test. **E)** Confocal images of DIV12-13 JIP3 KO i^3^Neuron somas pre-loaded with 25µg/mL DQ-Red BSA and treated with 0.15% DMSO or 15µM RH1115. Scale bar, 5µm. **F)** Quantification of the mean intensity of DQ-Red BSA. Superplots show mean ± SEM as well as individual data (represented as large and small circles, squares, and triangles). N=3 independent experiments, JIP3 KO DMSO n=64 i^3^Neurons, JIP3 KO RH1115 n=63 i^3^Neurons, \**p*<0.05, unpaired t-test.

We next examined the effect of RH1115 on the proteolytic activity of lysosomes in both Control and JIP3 KO i^3^Neurons using DQ-Red BSA (dye-quenched Bovine Serum Albumin), a cargo that fluoresces when proteolytically cleaved, in i^3^Neurons (Majumder et al., 2022; Marwaha & Sharma, 2017; Snead & Gowrishankar, 2022). To isolate the effect of RH1115 on lysosomal degradation alone and remove effects on DQ-Red BSA intensity arising from differences in endocytic uptake, lysosomes in both Control and JIP3 KO i^3^Neurons were preloaded with DQ-Red BSA prior to treatment with RH1115 or DMSO. The DQ-Red BSA intensity was decreased in both Control and JIP3 KO i^3^Neurons upon treatment with Bafilomycin A1 post lysosomal loading (Figure S2), confirming that it reports on lysosomal proteolytic activity. RH1115 treatment resulted in a striking increase in the DQ-Red BSA mean intensity in both Control and JIP3 KO i^3^Neurons, (Figure 3 C-F), suggestive of increased proteolytic activity in the lysosomes.

### The rescue of axonal lysosome pathology in JIP3 KO i^3^Neurons by RH1115 requires JIP4

Application of the small molecule RH1115, results in clearance of axonal lysosome accumulations in JIP3 KO i^3^Neurons and mobilizes lysosomes to the perinuclear region within the soma, suggesting an increase in retrograde lysosome transport. While JIP3 has an established role in regulating retrograde axonal lysosome movement (Drerup & Nechiporuk, 2013; Gowrishankar et al., 2017), the highly related adaptor JIP4 has some overlapping function in this pathway as well (Gowrishankar et al., 2021), and is known to play a role in dynein-dependent lysosome movement from the periphery to the perinuclear region in non-neuronal cells (Willett et al., 2017). The lysosomal transmembrane protein TMEM55B, has been shown to recruit JIP4 to lysosomes to induce this dynein-dependent movement in these cells (Willett et al., 2017). Given this prior evidence for regulation of TMEM55B expression as a means of regulating lysosome movement (Willett et al., 2017), we examined whether RH1115 treatment altered TMEM55B expression in these i^3^Neurons. Indeed, RH1115 treatment led to increased levels of TMEM55B protein in both Control and JIP3 KO i^3^Neurons (Figure 4 A, B). Considering the observed increase in TMEM55B levels and its known role in recruiting JIP4 to lysosomes to bring about retrograde movement, we interrogated if this rescue of axonal lysosome pathology in JIP3 KO i^3^Neurons depends on JIP4. For this, we examined the effect of RH1115 on the lysosomal pathology in JIP3/4 double KO i^3^Neurons stably expressing LAMP1-GFP. The LAMP1 accumulations, which are more severe in the JIP3/4 double KO i^3^Neurons (Gowrishankar et al., 2021) were not rescued by RH1115 treatment (Figure 4 C-E), suggesting that RH1115 requires JIP4 to mobilize the lysosomes and clear the axonal lysosome pathology, and thus likely enhances retrograde lysosome movement in neurons via a JIP4/TMEM55B-mediated pathway. Furthermore, the small molecule failed to mobilize lysosomes to the perinuclear region in the neuronal cell bodies of JIP3/4 double KO i^3^Neurons (Figure S3B) unlike what we observe in JIP3 KO i^3^Neurons (Figure S3A), suggesting a critical role for the adaptor JIP4 in retrograde lysosome movement in the neuronal cell body.

**Figure 4:**
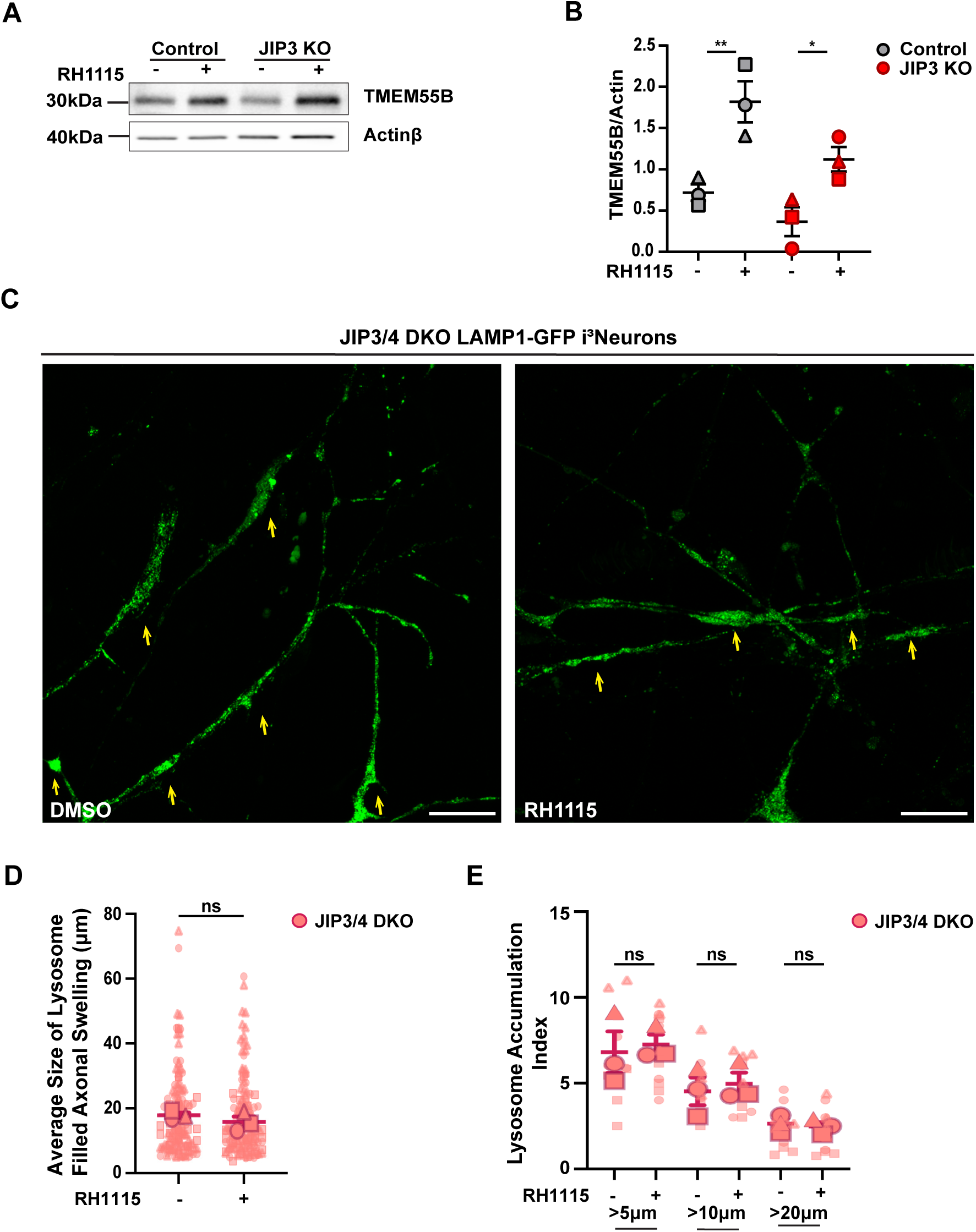
Net retrograde mobilization of axonal lysosomes induced by RH1115 in i^3^Neurons requires JIP3 or JIP4 and is accompanied by increased TMEM55B. **A)** Immunoblot showing TMEM55B expression in DIV21 Control and JIP3 KO i^3^Neurons treated with 0.15% DMSO or 15µM RH1115 for 72 h (Actinβ loading control). **B)** Quantification of TMEM55B levels normalized to the loading control. Plots show mean ± SEM N=3 independent experiments (represented as circles, triangles and squares); \**p*<.05, \*\**p*<.01; one-way ANOVA with Sidak’s multiple comparisons. **C)** High resolution, stitched images of DIV8 JIP3/4 DKO i^3^Neurons stably expressing LAMP1-GFP and treated with 0.15% DMSO or 15µM RH1115 for 72 h. LAMP1-positive vesicle accumulations are marked by yellow arrows. Scale bar, 20µm. **D)** Quantification of the length of LAMP1-positive vesicle accumulations in DIV8 JIP3/4 DKO LAMP1-GFP i^3^Neurons treated with 0.15% DMSO or 15µM RH1115. Superplots show mean ± SEM as well as individual data (represented as large and small circles, squares, and triangles). N=3 (here includes 2 independent experiments, and a technical replicate). DMSO treated n=61 swellings, RH1115 treated n=68 swellings. **E)** Quantification of the lysosome accumulation index (swellings greater than 5μm, 10μm and 20μm normalized to neurite density). Superplots show mean ± SEM as well as individual data (represented as large and small circles, squares, and triangles). N=3 (here includes 2 independent experiments, and a technical replicate), DMSO treated n=12 stitched images, RH1115 treated n**=12 stitched images.**

### RH1115 rescues locomotor defects in JIP3KO zebrafish larvae

Loss of JIP3 in zebrafish has been shown to alter retrograde axonal lysosome transport (Drerup & Nechiporuk, 2013). Our studies examining locomotion behavior in zebrafish larvae, revealed that JIP3 KO zebrafish move less distance and travel at slower velocity on average, compared to wild type clutch mates (Figure 5A, B). Treatment of JIP3 KO zebrafish with RH1115 for 6 days, immediately following fertilization, resulted in a striking rescue of the locomotor defects in fish that completely lack JIP3 protein, with significant increases in total distance travelled and average velocity (Figure 5A, B). In fact, the small molecule restored locomotor properties in the JIP3 KO zebrafish to levels observed in the wildtype zebrafish (Figure 5A, B), with no negative impacts on zebrafish larvae of either genotype.

**Figure 5:**
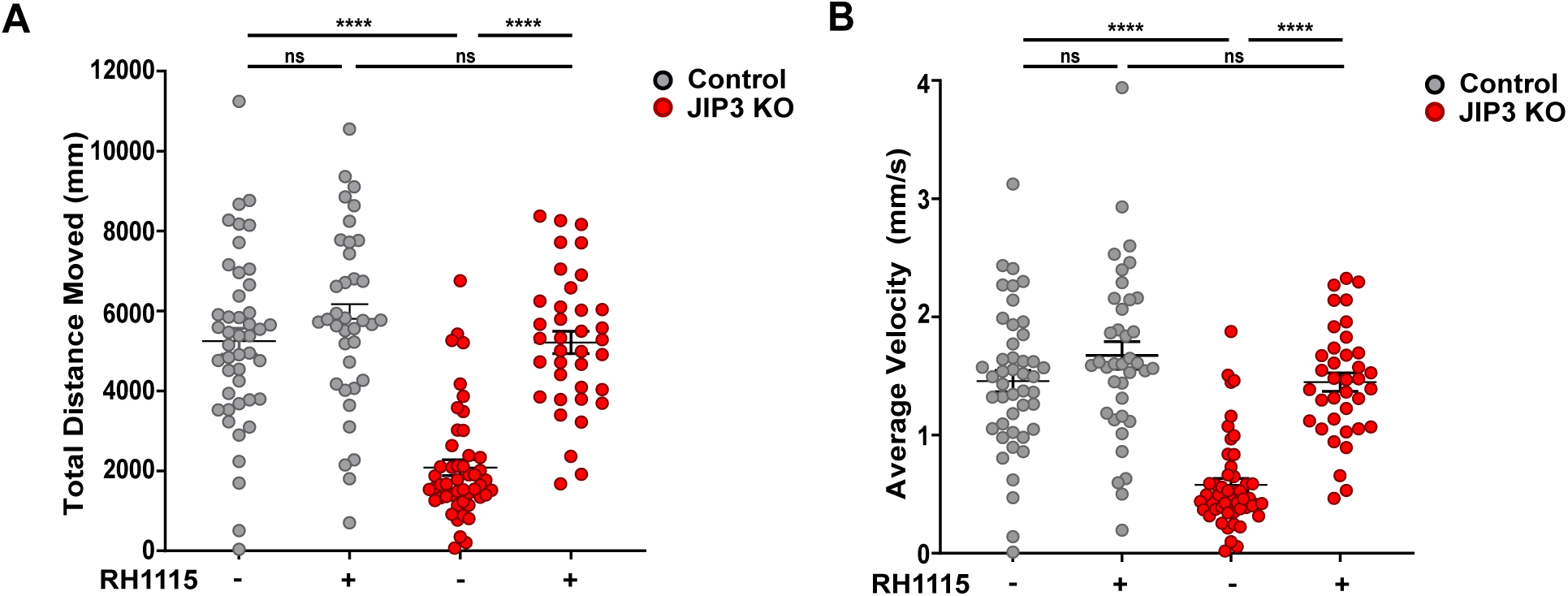
RH1115 treatment rescues locomotor defects in JIP3 KO larval zebrafish. **A)** and **B)** Decreased total distance moved and average velocity were displayed in JIP3 homozygous knockout (KO) zebrafish larvae, compared to JIP3 wild type (WT) clutch mates. Larval zebrafish treated with 0.05% DMSO or 0.15 μM RH1115 immediately following fertilization for 6 days were utilized for locomotor assay. At 6 days post-fertilization (dpf) total distance moved (mm) **A)** and average velocity (mm/s) **B)** of larval zebrafish were compared. Both locomotor phenotypes were shown to be rescued in JIP3 KO zebrafish treated with 0.15 μM RH1115 for 6 days. Each dot represents an individual 6-dpf larvae; WT DMSO N=45, WT RH1115 N=38, JIP3 KO DMSO N=49, JIP3 KO RH1115 N=37; ****p<0.0001, ns=non-significant, one-way ANOVA with Tukey’s test for multiple comparisons.

## Discussion

Perturbations to axonal lysosome transport that result in an imbalance in lysosomal distribution in axons has been linked to defects in neurodevelopment and to the pathophysiology of neurological diseases (Allison et al., 2017; De Pace, Ghosh, et al., 2024; De Pace, Maroofian, et al., 2024; Edmison et al., 2021; Farías et al., 2017; Gowrishankar et al., 2017; Paumier & Gowrishankar, 2024; Roney et al., 2021, 2022). The abnormal accumulation of protease-deficient axonal lysosomes near plaques (Gowrishankar et al., 2015), which were linked to APP processing, suggested that defects in retrograde transport and maturation of these organelles could contribute to plaque development. In support of such a model, loss of the brain-enriched lysosomal adaptor JIP3, recently shown to be a dynein activator (Cason & Holzbaur, 2023; Singh et al., 2024), resulted in axonal lysosome buildup as well as increased Aβ42 production in both primary mouse neurons and human iPSC-derived neurons (Gowrishankar et al., 2017, 2021). Furthermore, reducing dosage of JIP3 in a mouse model of AD resulted in increased soluble Aβ42 levels and increased neuritic plaque abundance and size (Gowrishankar et al., 2017). Together, these prior studies suggest that efficient retrograde axonal lysosome transport is anti-amyloidogenic and an attractive pathway for therapeutic intervention. In this study, we show that a small molecule modulator of ALP recently developed by our two groups, RH1115, clears axonal lysosome accumulation and reduces the extracellular Aβ42 levels in human i^3^Neurons lacking JIP3. RH1115 treatment clears not only the endolysosomal buildup, but also the autophagic vacuole accumulation from JIP3 KO axons, suggesting that it can restore protein and organelle homeostasis in neurons where axonal ALP is impaired. Thus, elucidating the mechanism of action for RH1115 in human neurons could be impactful in different neurodegenerative diseases where impaired ALP is a contributing factor.

Previous studies based on JIP3 depletion in PC12 cells and exogenous JIP3 expression in neurons postulated that JIP3 interacts with AVs marked by both LAMP1 and LC3, and that JIP3 modulates movement of more mature AVs in middle and proximal axons (Cason et al., 2021). Consistent with these reports (Cason et al., 2021; Cason & Holzbaur, 2023), our studies suggest that JIP3 affects the motility of autolysosomes (Figure 1 A-C). Here, we find that loss of JIP3 also impacts the motility of autophagosomes in addition to the movement of autolysosomes (Figure 1 A-C). Interestingly, several accessory proteins in addition to JIP3 have been identified to regulate the retrograde movement of AVs, including JIP1, HAP1-Huntingtin and RILP (Fu & Holzbaur, 2014; Hill et al., 2019; Khobrekar et al., 2020; Wong & Holzbaur, 2014). These adaptors have been suggested to act sequentially on AVs depending on the AV maturation state and spatial distribution within the axon (Cason et al., 2021). Our results from imaging stably expressed LC3-RFP-GFP in i^3^Neurons suggest that loss of JIP3 leads to a decrease in the motile fraction of both autophagosomes and autolysosomes but does not affect fusion of the axonal autophagosomes with endo-lysosomes. Given evidence from previous studies that the less mature autophagosomes are more frequent in the distal axon (Cason & Holzbaur, 2023; Cheng et al., 2015a; Maday et al., 2012), JIP3 likely has a role in regulating organelle transport in the distal axon as well. This is supported by endogenous JIP3 localization in distal axons, including in the tips of growth cones in neurons (Gowrishankar et al., 2017; Watt et al., 2015).

We also show that this rescue of axonal lysosome buildup in JIP3 KO i^3^Neurons requires the related lysosome adaptor JIP4 and that the small molecule RH1115 increases levels of the JIP4-interacting, transmembrane lysosomal protein TMEM55B in i^3^Neurons (Figure 4). TMEM55B has been shown to recruit JIP4 to lysosomes, and its upregulation linked to dynein-dependent lysosome movement in response to nutrient starvation in cultured cell lines (Willett et al., 2017). Thus, it is possible that RH1115-mediated TMEM55B upregulation recruits more JIP4 to axonal lysosomes and aids in their retrograde movement. A recent study demonstrated that TMEM55B interacts with an E3 Ubiquitin Ligase, NEDD4, in response to oxidative stress, which competes with the TMEM55B-JIP4 interaction (Jeong et al., 2024). It will be interesting to determine if RH1115 can modulate this NEDD4-TMEM55B interaction in response to oxidative stress. Of relevance to Parkinson’s disease, expression of a hyperactive LRRK2 in neurons led to increased JIP4 recruitment to AVs and also reduced their processive retrograde movement (Boecker et al., 2021). Our studies in the JIP3/4 DKO i^3^Neurons suggest that RH1115-mediated retrograde axonal lysosome movement requires JIP4. Given that prior studies implicated a JIP4-mediated bias towards kinesin activation on these AVs under pathological conditions, it may be possible that RH1115 could rescue the AV transport defect in the LRRK2 mutant neurons.

Our prior target identification studies revealed a specific interaction of RH1115 with LAMP1 (Hippman et al., 2023). LAMP1 is an abundant glycoprotein on the surface of late endosomes, degradative lysosomes, and biosynthetic precursor organelles (Cheng et al., 2018; Farías et al., 2017; Gowrishankar et al., 2015; Lie et al., 2021). While LAMP1 has been widely used as a ‘marker’ for this spectrum of organelles, its precise function in the context of lysosome transport is not fully understood. Based on the interaction with RH1115 and its effect on increasing net retrograde lysosome movement, it is tempting to speculate that LAMP1 may be involved in lysosome transport. Consistent with this possibility, LAMP1-APEX studies in neurons suggest that LAMP1 interacts with two critical GTPases involved in lysosome biogenesis and transport, namely ARL8B and Rab7 (Frankenfield et al., 2020). To our knowledge, RH1115 is the first compound targeting LAMP1 that alters lysosome movement. Future studies where we examine differential LAMP1 interactions upon RH1115 treatment, could shed light on how LAMP1 is involved in modulating lysosomal axonal transport.

RH1115 appears to enhance neuronal lysosome acidification (Figure 3). This change in lysosomal physiology may be mediated by its interaction with LAMP1 or its effect on TMEM55B. LAMP1 has been shown to interact with multiple subunits of the vacuolar ATPase via LAMP1-APEX proteomic analysis (Frankenfield et al., 2020). Previous research has shown that changes in nutrient status can modulate the association between the transmembrane V_0_ subunit and the membrane associated V_1_ subunit on the lysosomal membrane, determining lysosomal acidification (Ratto et al., 2022), thus it is possible that RH1115 may positively modulate vATPase subunit V_0_ or V_1_ interaction with LAMP1, increasing vATPase assembly. In addition, LAMP1 has also been shown to bind to and inhibit the proton leak channel in lysosomes, TMEM175, thus increasing acidification of lysosomes (Zhang et al., 2023). Alternatively, TMEM55B may also have a role in vATPase assembly on lysosomes that could contribute to observed effects of RH1115. TMEM55B deficiency in Hela cells causes a decrease in the levels of the ATP6V1A subunit of the vATPase in the detergent resistant membrane fraction (representing lipid rafts present in lysosomal membranes) suggesting that TMEM55B may have a role in anchoring or increasing the association of the V1 subunit with the lysosomal membrane (Hashimoto et al., 2018). Therefore, RH1115 increasing TMEM55B may increase lysosomal acidification through modulating vATPase association/assembly on lysosomal membranes. Future studies using the small molecule could also shed light on whether LAMP1 plays a role in modulating neuronal lysosomal acidification.

Additionally, the small molecule RH1115 increases perinuclear localization of lysosomes within the neuronal cell body/soma of JIP3 KO i^3^Neurons and increases lysosomal size, and lysosomal degradative capacity (Figure 2 E-H; Figure 3). Defects in lysosomal degradative capacity are a major contributing factor to the pathology of lysosomal storage disorders (Ballabio & Bonifacino, 2020; Boustany, 2013), therefore, effect of RH1115 on neuronal lysosomes could have a broad clinical relevance. In addition to rescuing autophagic and lysosomal defects in human neurons, RH1115 treatment rescues locomotor defects in JIP3 KO zebrafish larvae (Figure 5), suggesting that restoration of axonal lysosome transport can positively impact neuronal health and functioning.

The movement and positioning of lysosomes within the neuronal cell body could be influenced by a variety of cellular factors. For instance, in non-neuronal cells, lysosome positioning is altered by nutrient status (Korolchuk et al., 2011). In turn, the positioning of lysosomes affects cellular functioning including autophagy, cell migration, cancer cell invasion and antigen-presentation (Ballabio & Bonifacino, 2020). Future studies will determine how the small molecule could modulate these different processes in the distinct cell types.

In conclusion, our work identifies a small molecule modulator that enhances net retrograde lysosome movement, demonstrates that increasing efficient axonal lysosome transport can be anti-amyloidogenic, and provides a new molecular tool to both modulate lysosome position-dependent cellular functions and to elucidate the role of LAMP1 in regulating lysosome transport, acidification, and degradation.

## Methods

### iPSC Culture and i^3^Neuron Differentiation

The JIP3 KO and JIP3/4 DKO iPSC lines (generated from WTC-11 iPSC parental line) were described previously (Gowrishankar et al., 2021). iPSC cell lines were maintained in E8 media (Life Technologies) supplemented with .05% Penicillin/Streptomycin (Gibco) and were passaged when 70% confluent using Accutase (Corning). The iPSCs were differentiated into i^3^Neurons as described previously (Fernandopulle et al., 2018; Gowrishankar et al., 2021). i^3^Neurons were plated at 30,000 cells per 35 mm glass-bottom dishes (MatTek Life Sciences) for live imaging experiments or on 35mm glass coverslips (Carolina Biologicals) for immunofluorescence studies. Glass was pre-coated with 0.1 mg/ml Poly-L-Ornithine (Sigma Aldrich) and 10 μg/ml mouse Laminin (Gibco). I^3^Neurons were plated and maintained in Cortical Neuron Culture Medium containing KO DMEM F12 (Gibco) B27 supplement (Thermo Fisher), 10 ng/ml BDNF (PeproTech) and 10 ng/ml NT3 (PeproTech), 1 μg/ml mouse Laminin (Gibco), and 2 μg/ml Doxycycline (Fisher Bioreagents).

### Immunoblotting

Lysis of i^3^Neurons and western blotting was carried out as described previously (Gowrishankar et al., 2021). Briefly, DIV21 i^3^Neurons were washed three times with cold PBS and lysed in lysis buffer [1% Triton-X in PBS, Benzonase (Millipore Sigma, E1014), protease inhibitor (Thermo Fisher Scientific) and phosphatase inhibitor (PhosStop, Roche)]. Prior to immunoblotting, samples were run using SDS-PAGE for 1 hour and 20 minutes at 90V followed by transfer onto a nitrocellulose membrane. See Supplemental Table 1 for antibody information.

### Immunofluorescence analysis of i ^3^Neurons

i^3^Neurons differentiated for 1–2 weeks on 24 mm glass coverslips or 35 mm Mattek glass-bottom dishes were processed for immunostaining as described previously (Gowrishankar et al., 2017). See Supplemental Table 1 for antibody information.

### I^3^Neuron Viability Assay

DIV10 i^3^Neurons on 35mm glass coverslips were treated with 0.1% DMSO or 15μM RH1115 for 72 hours before fixation and immunostaining for Tau and LAMP1 as described previously. (Gowrishankar et al., 2021) Images were acquired (5 to 6 areas at random) using a high magnification objective on the Keyence BZ-X810 microscope (Osaka, Japan), and the number of neurons per unit area was computed. Tau staining was used to confirm neuronal viability (normal morphology and neurite integrity). Mean ± SEM of three independent experiments was computed.

### Analysis of Autophagosome and Autolysosome Distribution and Motility

DIV15-16 i^3^Neurons stably expressing LC3-RFP-GFP were imaged live at 37°C for approximately 20 minutes. Neuronal processes were selected at random after ensuring that they were sufficiently dispersed to allow for easy identification of individual processes. Time-lapse images in both green and red channels were acquired at 2.5 FPS using Fast Airyscan mode on LSM880 microscope (Zeiss) with a 63× oil objective (1.4 NA) and 2.5× optical zoom. Motile fraction was analyzed as described previously (Gowrishankar, JCB, 2017). RFP and GFP intensities were scaled based on control condition and then autophagosomes (Green + Red) and autolysosomes (Red) were identified and tracked. Every neurite (5-17 per time series) of at least 10µm was analyzed. Neurites were traced using the ‘freehand line’ tool, followed by straightening using ‘straighten’ tool, and then ‘resliced’ to create a kymograph (ImageJ). Vesicles were identified as autolysosomes if they did not contain any green channel puncta. Vesicles were identified as motile if they moved 2µm or more during the period of the movie (2 minutes). Vesicle density was quantified from a single timeframe (t=10) for all neuronal processes examined. % Autolysosomes (RFP only) per neuronal process was also computed from timeframe t=10.

### Quantification of Axonal Lysosome Accumulation Index

Axonal lysosome buildup was quantified using endogenous LAMP1 staining or from LAMP1-GFP fluorescence. For experiments involving analysis of endogenous LAMP1 staining, DIV10-12 Control and JIP3 KO i^3^Neurons were treated with 0.15% DMSO or 15μM RH1115 for 72 hours and fixed and stained for LAMP1. For exogenous LAMP1-GFP experiments, DIV10-14 JIP3 KO or DIV 8 JIP3/JIP4 DKO LAMP1-GFP i^3^Neurons were fixed and imaged. 3×3 stitched z-stack images were acquired using Airyscan SR (super resolution) imaging mode on an LSM880 Confocal microscope (Zeiss) with a 63× oil immersion objection (1.4 NA) to capture 200μm^2^ area per image, at high resolution. Stitched images were analyzed by setting the ‘threshold’ for visualizing LAMP1+ swellings based on the DMSO condition in JIP3 KO i^3^Neurons. All LAMP1+ swellings above that threshold were counted using the ‘line’ tool followed by the ‘measure’ tool. The lysosome accumulation index was determined by counting the number of swellings greater than 5µm, 10µm, and 20µm (for JIP3/4 DKO only) per 200µm^2^ image area and then normalizing to the number of neurites in that area, this number was then multiplied by 10. The lysosome accumulation index is normalized to the mean of the control condition.

### Analysis of Lysosomal vesicle Properties in i^3^Neurons

ImageJ ‘analyze particles’ tool was used to measure lysosomal vesicle size, integrated density, and number within i^3^Neuron somas immunostained for LAMP1. Each soma was outlined using the ‘freehand selections’ tool, and then using ‘threshold’ function, a threshold was set based on optimal coverage of most vesicles possible without compromising vesicle size and shape (avoiding false negatives or collapse/fusion of different vesicles). The ‘analyze particles’ tool was then used to define the parameters for the measured vesicles, with a minimum size of 0.002 µm^2^ and no maximum size. Mean size, intensity of vesicles, and count per neuron were computed.

### Evaluation of Lysosome Positioning in JIP3 KO i^3^Neurons

DIV10 i^3^Neurons on 35 mm glass coverslips treated with 0.1% DMSO, or RH1115 (15 μM) for 72 hours were fixed and stained for Tau, LAMP1 and DAPI as described previously (Hippman et al., 2023). Images were acquired using a 60X objective on the Keyence BZ-X810 microscope. Lysosome distribution was classified as “clustered”, “normal”, or “intermediate” based on proximity to nucleus (DAPI signal). Imaging and analysis were carried out in a double-blind fashion. Mean ± SEM of percent neurons exhibiting perinuclear clustering of lysosomes from three independent experiments were computed.

### Evaluation of Lysosome Positioning in Control Hela cells expressing LAMP1-KBS-GFP

Control HeLa cells were transfected with LAMP1-KBS-GFP plasmid (Pu et al., 2015) using Fugene reagent (Promega) for 48 hours and then treated with 0.15% DMSO or 15μM RH1115 for 6 hours, fixed, and stained with anti-GFP (Invitrogen) antibody and DAPI (SouthernBiotech). Images were acquired on an LSM880 in confocal mode using 63× oil immersion objective at 1.5× optical zoom. Cells of comparable GFP intensities, and in smaller clusters of 2-5 cells were chosen to ensure uniformity in morphology. Cells were categorized into three groups, blinded to condition, based on where the majority of LAMP1-GFP signal was seen within the cell: majority seen in periphery, majority seen in perinuclear region, or mixed distribution (equally dispersed or equal intensity in periphery and perinuclear).

### Evaluation of Lysosome Acidification in i^3^Neurons

DIV10-13 Control and JIP3 KO i^3^Neurons were incubated with 100nM of LysoTracker-Red (Supplemental Table 2) for 10 minutes in culture medium at 37°C. After incubation, i^3^Neurons were washed in warm imaging media twice before imaging live for approximately 15 minutes per condition at 37°C using an LSM880 microscope in Airyscan super resolution mode using a 63× oil immersion objective (Zeiss). Images were acquired after focusing on central plane of the soma. Images were processed using Airyscan processing function of Zen software. Lysotracker intensity was measured in ImageJ using the ‘freehand’ selection tool to outline the soma followed by the ‘measure’ tool. Mean intensity per cell was computed and population mean across multiple independent experiments across treatments were compared.

### Lysosomal Degradation Assay

Lysosomal degradation was measured using DQ-Red BSA (Supplemental Table 2) as described previously (Snead & Gowrishankar, 2022) with some modifications. I^3^Neurons were preloaded with DQ-BSA prior to treatment with 0.15% DMSO or 15μM RH1115 (Ratto et al., 2022). Conditioned media from DIV12-13 i^3^Neurons was saved and DQ-Red BSA (25µg/mL) that was equilibrated in 10% culture media at 37°C for 5 minutes was added to the i^3^Neurons and incubated for 4 hours at 37°C, 5% CO_2_. DQ-Red BSA solution was removed and i^3^Neurons were washed with warm PBS twice before a 2-hour chase in the conditioned media. Conditioned media was replaced with 10% culture media containing 0.15% DMSO or 15μM RH1115 for 4.5 hours. i^3^Neurons were washed twice with warm imaging media (20 mM HEPES, 5 mM KCL, 1 mM CaCl2, 150 mM NaCl, 1 mM MgCl2, 1.9 mg/mL glucose and BSA, pH 7.4) prior to imaging live at 37°C on an LSM880 in Airyscan super resolution mode, on a 63× oil immersion objective, focusing on the central plane of the somas. Images were processed using ‘Airyscan processing’ function of Zen software and the intensity was analyzed in ImageJ using the ‘freehand’ selection tool to outline the soma followed by the ‘measure’ tool.

### Measurement of Extracellular Aβ42

Control and JIP3 KO i^3^Neurons were differentiated for 42 days. At DIV42 the media was removed and replaced with KO DMEMF12 (equilibrated overnight at 37°C and 5% CO_2_) with DMSO (0.15%), BACE1-I (5μM) or small molecule (RH1115 15μM). i^3^Neurons were maintained for 7 days in KO DMEMF12 + treatment, with a media change and addition of new DMSO/BACE1-I/RH1115 4 days after the first treatment (Fig 2 C, D). On DIV49/50 i^3^Neurons were lysed and conditioned media was spun down at 5000g for 5 min and the supernatant saved at -80°C. CM was measured using ELISA Aβ42 high sensitivity kit (Wako) according to the manufacturer’s instructions.

### Generation and Maintenance of Zebrafish lines

A previously described JIP3 knockout (KO) zebrafish line (JIP3^n17^) was obtained (Drerup & Nechiporuk, 2013). Briefly, JIP3 KO fish were generated using ENU mutagenesis and found to have a single nucleotide substitution that resulted in an early stop codon, SpeI restriction enzyme site, and truncated JIP3 protein. Genotyping was conducted using PCR with the following primers: Forward – 5’-TTTGTCTGTTGAAATTGCT-3; Reverse – 5’-ACGGTCCATACCCATGATT-3’, to generate a 385-base pair (bp) amplicon followed by an overnight restriction enzyme digest with SpeI (New England Biosystems, Ipswich MA) to generate double band homozygous amplicons (243 and 142 bp) detected using gel electrophoresis.

Animals were raised in quarantine and housed at the University of Alabama at Birmingham (UAB) Zebrafish Research Facility and all procedures were approved by the UAB Institutional Animal Care and Use Committee. Zebrafish embryos were raised in a 28.5°C incubator on a 14-hour light, 10-hour dark cycle to 5-days post-fertilization in 1 x E3B made in recirculating, filtered system water. After 5 days, animals were transferred to tanks on the recirculating water system (Aquaneering, Inc., San Diego, CA). Heterozygous JIP3 adult fish were in crossed (1 male:1 female) to generate wild type, heterozygous, and homozygous embryos.

### Zebrafish Locomotion Assay

Embryos were placed in petri dishes containing either 0.05% DMSO dissolved in E3 blue (no drug controls) or 0.05% DMSO with 0.15 μM RH1115 and kept in the incubator with steady temperature and light cycles. Water changes were completed daily in the morning to refresh drug and maintain a steady concentration during screening. At 6-days post fertilization larvae were transferred to 4 x 48-well plates, allowed to acclimate, and then run on the Zantiks (Zantiks Ltd, Cambridge UK) for 1 hour to observe locomotor activity. Using a camera and infrared light tracing, total distance travelled (mm) and average velocity (mm/s) were calculated for individual larvae. Larvae were then transferred to a PCR genotyping plate and DNA was extracted using NaOH, neutralized with TRIS-HCL, and genotyped as described above. One-way ANOVA with Tukey’s test for multiple comparisons was used to calculate significance across genotypes for locomotor phenotypes and all analyses were completed using GraphPad Prism Software (GraphPad Software, LLC, San Diego, CA).

### Statistical Analysis

Data are represented as superplots consisting of mean ± SEM across experiments as well as individual data points across all the experiments, unless otherwise specified. Statistical analysis was performed using GraphPad Prism 10 software. Groups (means from the different experimental repeats) were compared using students t-test or one-way ANOVA multiple comparison test as appropriate. Detailed statistical information [number of independent experiments(N), number of individual measurements(n), statistical test performed and *p* values] are described in the respective figure legends.

## Acknowledgement

We thank Shawn Ferguson, Yale University for generously sharing the JIP3/4 DKO iPSC line. We thank Dr. Alex Nechiporuk and his team at Oregon Health and Sciences University for providing us with their JIP3 KO zebrafish line for our behavioral phenotyping efforts.

## Author Contributions

DD, NC equal contributions. RS, RL equal contributions. AMS, SP, MK, GS, CC, SG designed experiments. SMC and LNA contributed to designing experiments. AMS, SP, MK, RH, GS, RSH, DD, NC, RS, RL, AD, CC, SG performed experiments. RSH, AD carried out synthesis of small molecule. AMS, SP, MK, DD, NC, RS, RL, SG performed experiments in i^3^Neurons. GS and CC performed experiments in zebrafish. AMS, SP, MK, GS, DD, NC, RL, CC, SG analyzed data. AMS, MK, GS prepared figures. AMS and SG prepared the initial draft of the manuscript with help from MK. SMC and LNA contributed to editing the manuscript. LNA directed and supervised chemical synthesis. SG, LNA and CC secured funding for the project. SG directed the project and led the efforts to prepare the manuscript.

## Funding

Research reported in this publication was supported by the Wolverine Foundation (SG, CC), and the National Institute on Aging and National Cancer Institute of the National Institutes of Health [RF1AG076653 (SG), R01AG074248 (SG), T32AG057468 (AMS), F30AG081091 (AMS) and F31CA265072(RSH)]. This research was also supported in part by the intramural program of the National Institutes of Health. The content is solely the responsibility of the authors and does not necessarily represent the official views of the National Institutes of Health.

## Conflict of Interest

LNA, RSH and AD are named as inventors on the patent of the intellectual property on RH1115 owned by UIC and the U.S government. The other authors declare that they have no competing financial interest.

