## Supplemental Figures and Legend for "Small molecule modulator of neuronal lysosome positioning and function resolves Alzheimer’s Disease-linked pathologies in cultured human neurons"

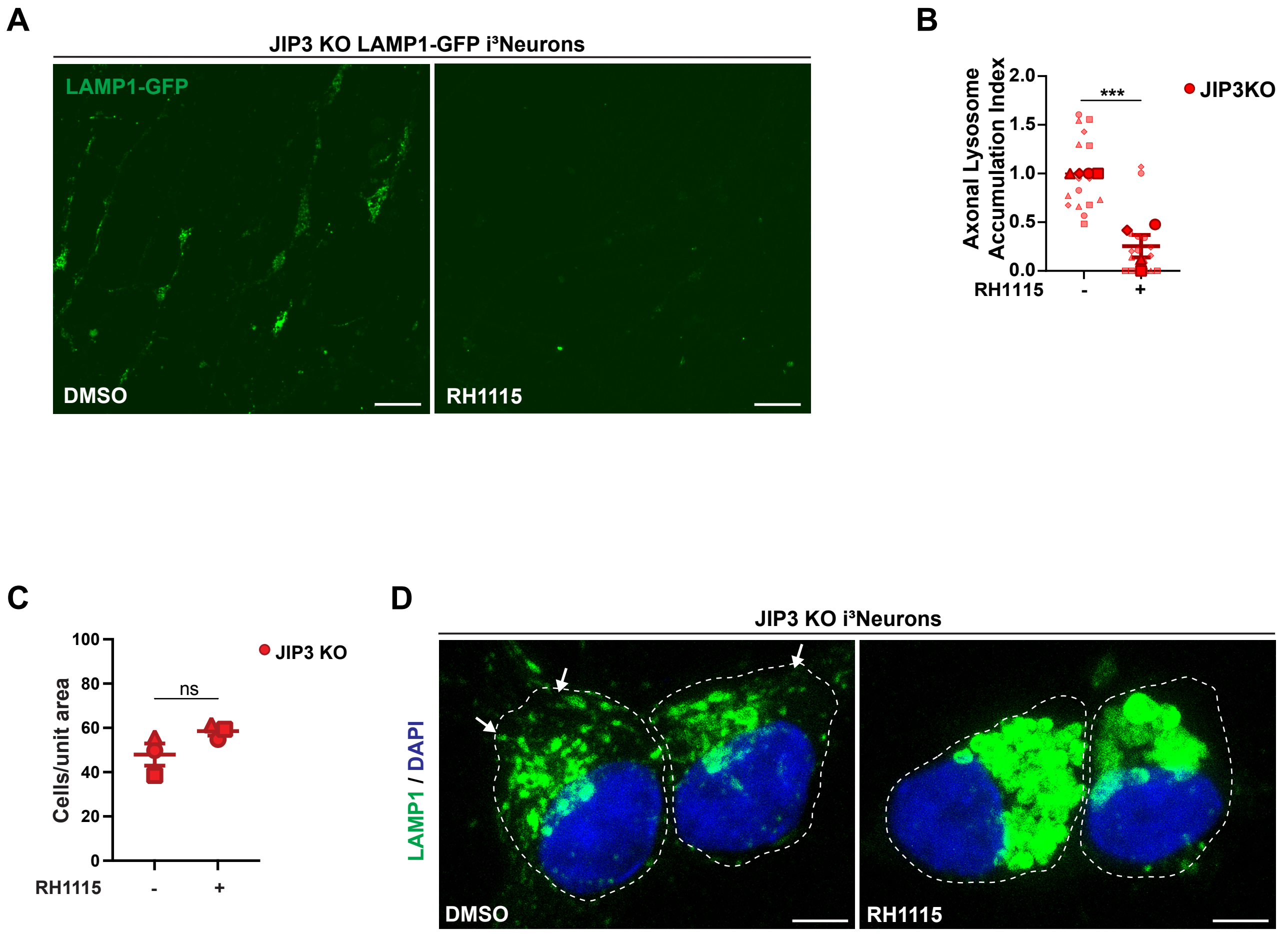
A
JIP3 KO LAMP1-GFP i³Neurons
LAMP1-GFP
DMSO
RH1115
B
Axonal Lysosome Accumulation Index
2.0
1.5
1.0
0.5
0.0
***
JIP3KO
RH1115
-
+
C
Cells/unit area
100
80
60
40
20
0
ns
JIP3 KO
RH1115
-
+
D
JIP3 KO i³Neurons
LAMP1 / DAPI
DMSO
RH1115

**Figure S1**

**Supplemental Figure S1**: **RH1115 mobilizes LAMP1 vesicles in JIP3 KO i$^3$Neurons without reducing cell viability.**

**A)** Representative images of DIV10 JIP3 KO LAMP1-GFP i$^3$Neurons treated with either 0.15% DMSO or 15µM RH1115 for 72 h. Scale bar, 10µm. **B)** Quantification of the axonal lysosome accumulation index (swellings greater than 5µm normalized to neurite density). Superplots show normalized mean ± SEM as well as individual data (represented as large and small circles, triangles and squares). N=4 independent experiments, ***p<0.001; unpaired t-test. **C)** Quantification of the number of DIV10 JIP3 KO i$^3$Neurons per unit area, following treatment with 0.15% DMSO or 15µM RH1115 for 72 h, as a read out of neuronal viability. Data show mean ± SEM from N=3 independent experiments, n=~250 i$^3$Neurons per treatment per experiment; ns=non-significant. **D)** Representative images of DIV11 JIP3 KO i$^3$Neurons treated with either 0.15% DMSO or 15µM RH1115 for 72 h and stained for endogenous LAMP1 and DAPI. White dashed line outlines neuronal cell bodies. White arrowheads indicate peripheral lysosomes present in Control i$^3$Neurons. Brightness levels of LAMP1 vesicles in both images enhanced to allow visualization of dimmer peripheral lysosomes in the cell bodies. Scale bar, 5µm.

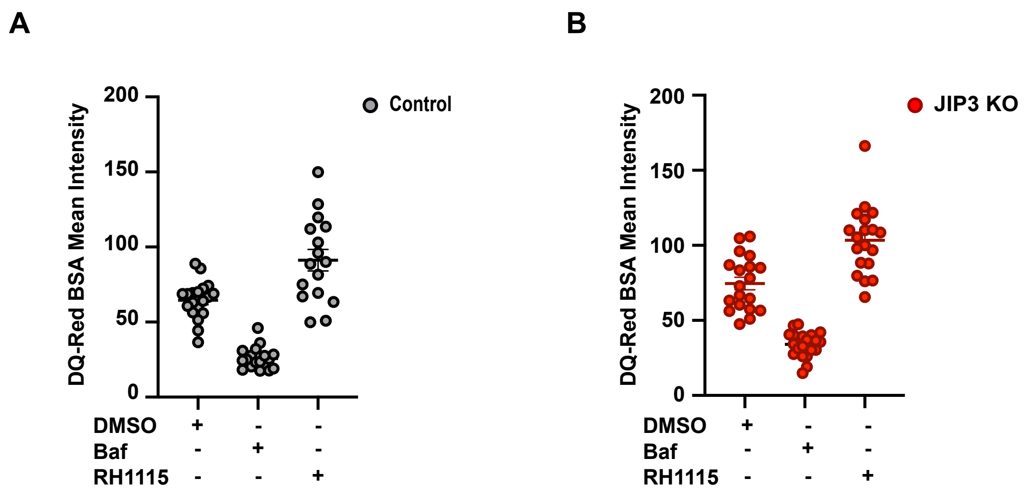
A
B
DQ-Red BSA Mean Intensity
200
150
100
50
0
Control
JIP3 KO
DMSO
Baf
RH1115
+
-

**Figure S2**

**Supplemental Figure S2: DQ-Red BSA  luorescence intensity is reduced with Ba ilomycin A1 treatment.**

**A)** Quantification of DQ-Red BSA fluorescence intensity (arbitrary units) in DIV12 Control $i^3$Neuron cell bodies. Cells were loaded with 25 µg/mL DQ-Red BSA probe and chased prior to 4.5 h treatment with 0.15% DMSO, 100nM Bafilomycin A1, or 15µM RH1115. Data from 1 independent experiment. DMSO=19, Baf=18, RH1115=16. **B)** Quantification of DQ-Red BSA fluorescence intensity (arbitrary units) in DIV12 JIP3 KO $i^3$Neuron cell bodies. DQ-Red BSA probe was loaded and chased prior to 4.5 h treatment with 0.15% DMSO, 100nM Bafilomycin A1, or 15µM RH1115. Data from 1 independent experiment. DMSO=19, Baf=21, RH1115=19.

A

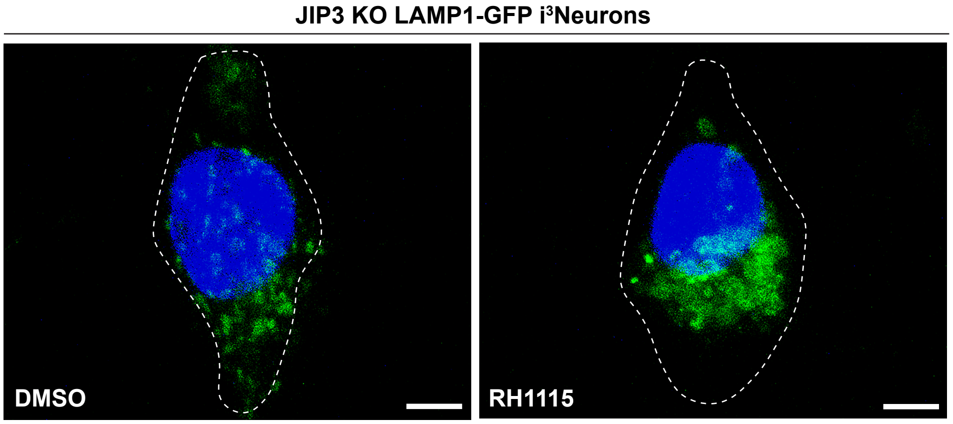
JIP3 KO LAMP1-GFP i$^3$Neurons
DMSO
RH1115

B

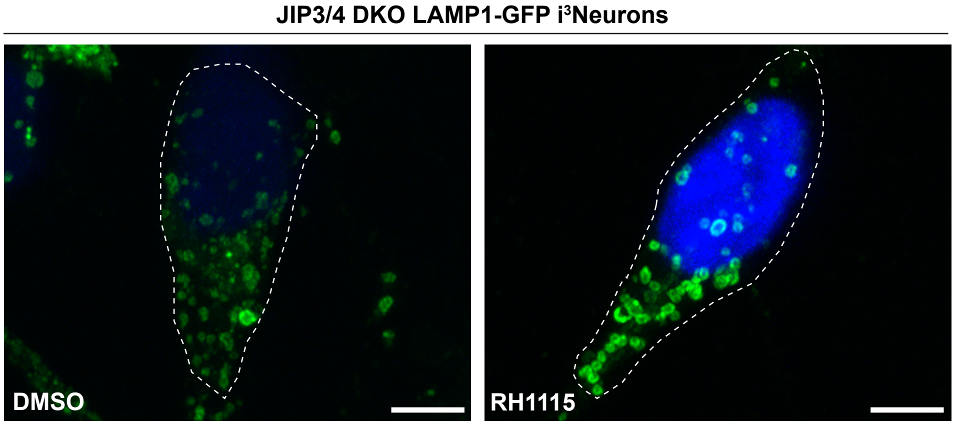
JIP3/4 DKO LAMP1-GFP i$^3$Neurons
DMSO
RH1115

Figure S3

**Supplemental Figure S3: LAMP1 distribution in soma of JIP3 KO and JIP3/4 DKO i$^3$Neurons.**

**A)** Representative high-resolution confocal images of soma of DIV8 JIP3 KO LAMP1-GFP i$^3$Neurons treated with either 0.15% DMSO or 15µM RH1115 for 72 h. Nuclei are stained with DAPI, somas are outlined. Scale bar, 4µm. **B)** Representative high-resolution confocal images of soma of DIV8 JIP3/4 DKO LAMP1-GFP i$^3$Neurons treated with either 0.15% DMSO or 15µM RH1115 for 72 h. Nuclei are stained with DAPI, somas are outlined. Scale bar, 4µm.

**Supplemental Table 1: Antibodies used in this study.**

| Antibodies | Source | Identifier |
|---|---|---|
| anti-ACTB Monoclonal | AB Clonal | AC026 |
| anti-hLAMP-1 Monoclonal | DSHB | H4A3 |
| anti-hLAMP-1 Monoclonal | Cell Signaling | 9091 |
| anti-LC3A/B Polyclonal | CST | 4108S |
| anti-MAP2B Monoclonal | BD Transduction Laboratories | 610460 |
| anti-TMEM55B Polyclonal | Invitrogen | PA5-61760 |
| anti-α-Tubulin Monoclonal | Sigma-Aldrich | T5168 |
| anti-mouse 488 | Thermo Fisher | A21202 |
| anti-rabbit 488 | Thermo Fisher | A21206 |
| anti-mouse 594 | Invitrogen | A21203 |
| anti-rabbit 594 | Invitrogen | A21207 |
| anti-rabbit HRP-linked antibody | CST | 7074 S |
| anti-mouse HRP-linked antibody | CST | 7076 S |
| anti-biotin HRP Linked antibody | CST | 7075 P5 |
| anti-GFP m3E6 Monoclonal | Invitrogen | A11120 |

**Supplemental Table 2: Reagents used in this study.**

| Reagents | Source | Identifier |
|---|---|---|
| Molecular Probes LysoTracker Red DND-99 | Invitrogen | L7528 |
| DQ Red BSA | Thermo Fisher | D12051 |
| DAPI Fluoromount-G | SouthernBiotech | 0100-20 |
